# Prion diseases disrupt the glutamate/glutamine metabolism in skeletal muscle

**DOI:** 10.1101/2023.10.31.564879

**Authors:** Davide Caredio, Maruša Koderman, Karl Frontzek, Silvia Sorce, Mario Nuvolone, Juliane Bremer, Petra Schwarz, Stefano Sellitto, Nathalie Streichenberger, Claudia Scheckel, Adriano Aguzzi

## Abstract

In prion diseases, aggregates of misfolded prion protein (PrP^Sc^) accumulate not only in the brain but also in extraneural organs. This raises the question whether prion-specific pathologies arise also extraneurally. Here we sequenced mRNA transcripts in skeletal muscle, spleen and blood of prion-inoculated mice at eight timepoints during disease progression. We detected gene-expression changes in all three organs, with skeletal muscle showing the most consistent alterations. The glutamate-ammonia ligase (*GLUL*) gene exhibited uniform upregulation in skeletal muscles of mice infected with three distinct scrapie prion strains (RML, ME7, and 22L) and in victims of human sporadic Creutzfeldt-Jakob disease. *GLUL* dysregulation was accompanied by changes in glutamate/glutamine metabolism, leading to reduced glutamate levels in skeletal muscle. None of these changes were observed in skeletal muscle of humans with amyotrophic lateral sclerosis, Alzheimer’s disease, or dementia with Lewy bodies, suggesting that they are specific to prion diseases. These findings reveal an unexpected metabolic dimension of prion infections and point to a potential role for GLUL dysregulation in the glutamate/glutamine metabolism in prion-affected skeletal muscle.

## Introduction

Prions are infectious protein aggregates that cause neurodegenerative diseases of the central nervous system (CNS). Prions multiplicate through the seeded conversion of the physiological cellular prion protein PrP^C^ into a misfolded, aggregated conformer termed PrP^Sc^ (1). PrP^C^ is expressed not only in the nervous system but also in the skeletal muscle and, to a lesser extent, in lymphoreticular tissue and blood (2, 3). Prions can be present in the blood, where they bind to plasminogen (4). Blood is a documented route of infection and remains a challenge for transfusion medicine (5, 6). Prions can enter the body through the gastrointestinal system and accumulate in lymphoid tissue, leading to neuroinvasion via peripheral nerves (7, 8). In variant CJD (vCJD), PrP^Sc^ is prominent in spleen, muscle, retina, blood vessels, skin, liver, kidney, and pancreas (9). Skeletal muscles of patients with acquired and sporadic CJD show PrP^Sc^ deposits in peripheral nerve fibers (10). Thus, while the most visible toxicity of prion diseases (PrDs) occurs in the brain, there is increasing evidence of peripheral manifestations which may be relevant to disease symptoms.

Growing evidence indicates that gene-expression changes in extraneural tissues, including blood, spleen, and skeletal muscle, can serve as markers of neurodegenerative disease progression (11–14). RNA sequencing of whole blood in Parkinson’s disease uncovered early immune cell changes and distinct gene expression patterns (15). A recent study linked exaggerated type I myofiber grouping in Parkinson’s Disease (PD) to altered gene expression in muscle, suggesting significant neuromuscular junction involvement and remodeling (16). Analogously, differential expression of muscle-specific genes was found in amyotrophic lateral sclerosis (ALS) patients, suggesting muscle-level changes alongside neural degeneration (17). Lymphoid tissue also accumulates prions (18, 19) and may experience molecular changes with diagnostic and prognostic potential.

Here we have conducted transcriptome-wide RNA sequencing analyses on blood, skeletal muscle, and spleen of mice after intracerebral exposure to prions and in autoptic skeletal muscle of humans diagnosed with sporadic CJD. We found that glutamate-ammonia ligase (*GLUL*) is uniquely upregulated in skeletal muscle of prion-infected mice and humans, but not in amyotrophic lateral sclerosis (ALS), Alzheimer’s disease (AD), or dementia with Lewy bodies (DLB). The pronounced increase of GLUL in prion-infected skeletal muscle, a distinctive feature not seen in ALS, AD, or DLB, underscores a unique aspect of prion diseases. This finding, in conjunction with observed reductions in glutamate levels in both animal models and human cases of prion infection, suggests a disruption in glutamate/glutamine metabolism in skeletal muscles as the disease progresses. This points to a prion-specific muscular pathophysiology diverging from other neurodegenerative disorders.

## Results

### Transcriptional derangement in skeletal muscle during prion disease progression

For this study, we used a previously established cohort of wild-type 2-month old C57BL/6 mice (20) which we had injected intracerebrally (i.c.) with scrapie prions (6^th^ consecutive mouse-to-mouse passage of mouse-adapted Rocky Mountain Laboratory sheep scrapie prions, abbreviated as RML6). For control, we injected non-infectious brain homogenate (NBH). Spleen, hindlimb skeletal muscle and blood were collected during necropsy at 4, 8, 12, 14, 16, 18 and 20 weeks-post-inoculation (wpi) as well as at the terminal stage of disease (Fig. 1A). We stratified our collective into three categories: early stage (4 and 8 wpi), pre-symptomatic stage (12, 14, 16 wpi) and symptomatic stage (18, 20, wpi and terminal) (21) (Fig. 1A).

**Fig. 1.**
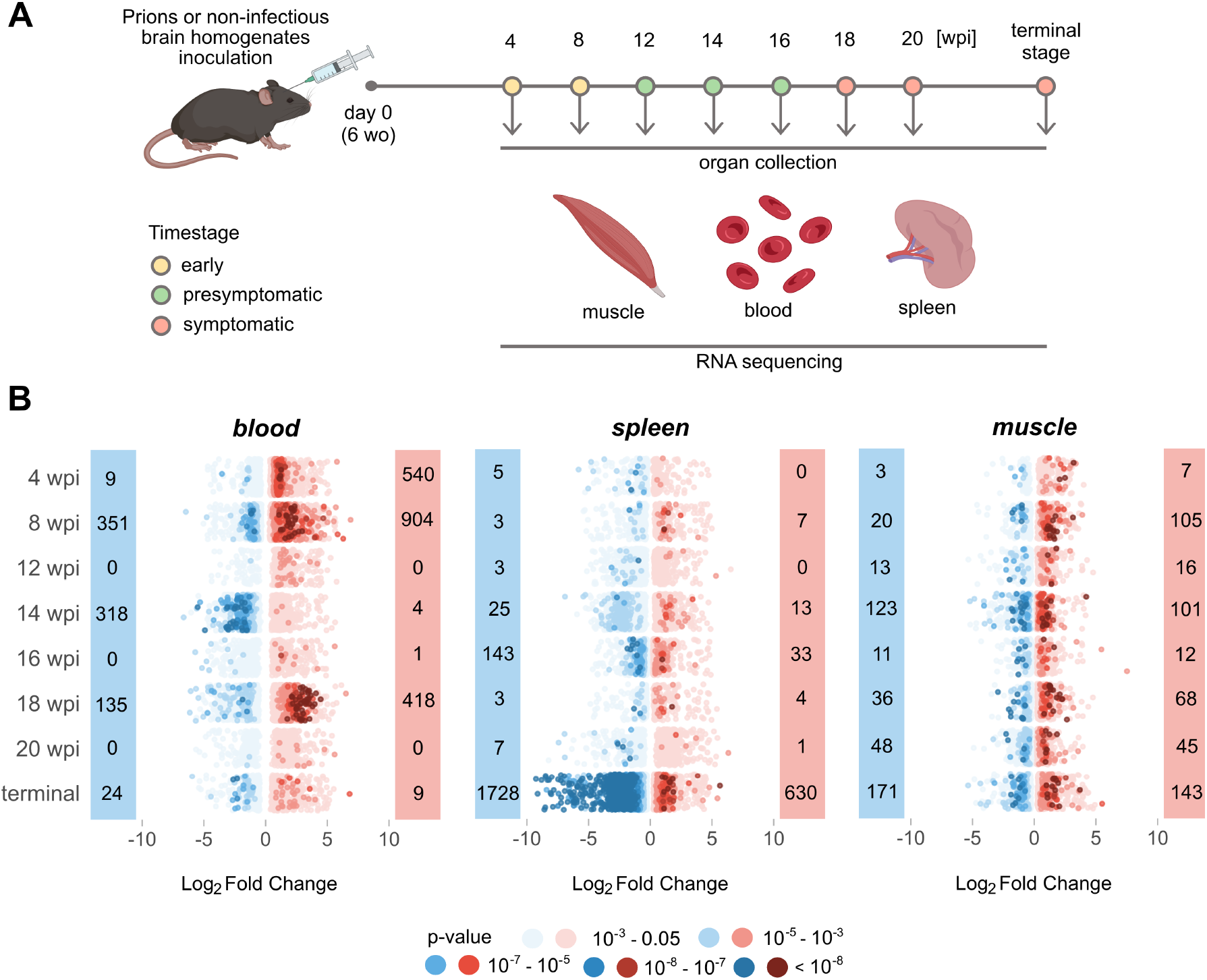
Temporal Dynamics of Gene Expression in Prion-Affected Tissues. **(A)** Muscle, blood, and spleen tissues were collected for bulk RNA sequencing at eight individual timepoints (wo = weeks old; wpi = week post inoculation). Samples were stratified into early, presymptomatic, and symptomatic stages. **(B)** Prevalence of upregulated (red) and downregulated (blue) DEGs (p-value < 0.05) across disease progression in the three tissues analysed. The dots in the dot plot represent individual genes and are color-coded according to their corresponding p-values.

We defined differentially expressed genes (DEGs) as transcripts with absolute log_2_ fold change |log_2_FC| > 0.5 and p-value < 0.05 (Fig. 1B). Transcriptional changes in blood were inhomogeneous during disease progression (Fig. 1B), possibly because peripheral blood may undergo changes in its cellular composition during infection or inflammation. Nevertheless, in both early timepoints DEGs associated with blood coagulation and hemostasis were detected (Supplementary Fig. 1, A and B; Supplementary Table 1-2). During the presymptomatic and symptomatic stages we did not identify any overlapping changes in blood. In contrast, major transcriptional changes in the spleen were found in terminal disease (Fig. 1B). Except for *Pcdh18*, which was significantly altered in both early stage timepoints, there was no overlap between DEGs in blood and spleen (Supplementary Table 3).

Compared to other analyzed organs, the number of DEGs in skeletal muscle remained relatively constant throughout the course of the disease (Fig. 1B). However, this consistent pattern was punctuated by recurrent up- or downregulation of specific genes at distinct disease stages. During the presymptomatic stage a single gene, *Adh1*, displayed upregulation, whereas the symptomatic stage featured elevated expression of *Mir8114*, *Glul*, and *Pik3r1* (Supplementary Table 3). To allow for interactive exploration of the results described in this study and for integration with our previously reported findings (20), we constructed a searchable database of gene expression profiles from brain and extraneural organs available for visualization and download at https://fgcz-shiny.uzh.ch/priontranscriptomics/.

### Consistently altered gene modules in skeletal muscle during prion disease progression

WGCNA (Weighted Gene Coexpression Network Analysis) identifies modules of highly correlated genes, which helps detecting coordinated changes in gene expression. We utilized WGCNA in conjunction with differential expression (DE) analysis to deduce organ-specific gene co-expression networks (Supplementary Table 4). To summarize the gene-expression levels of individual network modules, we calculated module eigengenes (MEs) representing the first principal component of each module. We identified 25 and 13 modules in blood and spleen, respectively, but we did not find any significant differences between the MEs of these modules in the two study groups across all three disease time stages (Supplementary Fig. 2, A and B). Conversely, in the muscle co-expression network, two of 39 modules (“orange” and “darkgreen”) showed significant differences in MEs between NBH controls and prions throughout disease progression (Supplementary Fig. 2C). The “orange” module (163 genes) was upregulated, while the “darkgreen” module (198 genes) was downregulated as the disease advanced (Fig. 2A).

**Fig. 2.**
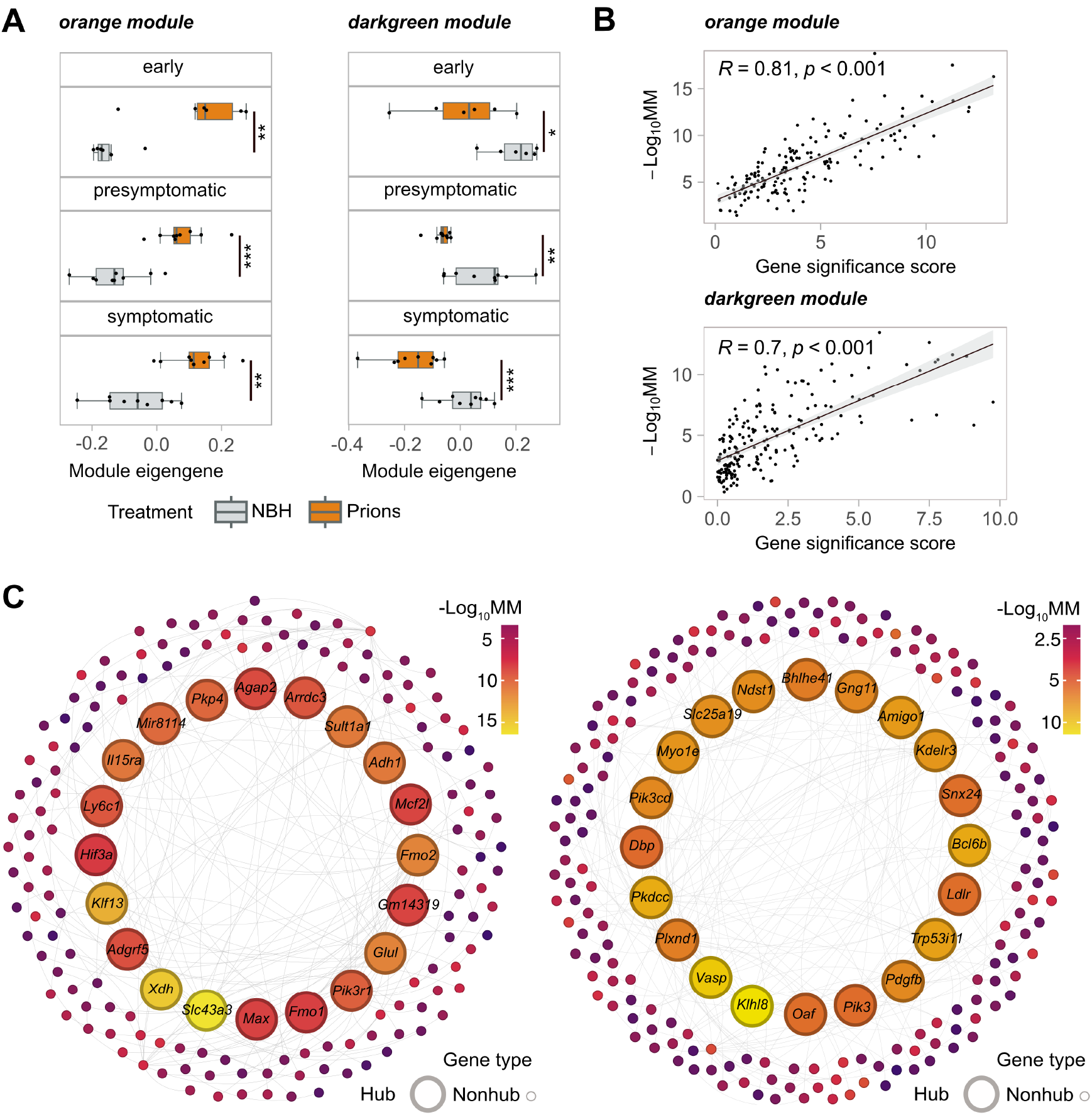
WGCNA Analysis of Gene Co-expression Modules. **(A)** Boxplots of module eigengenes of the main cohort for gene co-expression orange and darkgreen modules identified by WGCNA at different timestages (early, presymptomatic and symptomatic). Statistical significance (*p < 0.05, **p < 0.01, ***p < 0.005, ****p < 0.001) is indicated by asterisks **(B)** The scatter plots illustrate the relationship between the gene significance score and module membership (MM). Pearson correlation coefficient (R) and its corresponding p-value are displayed. **(C)** The minimum spanning trees with nodes representing genes within the orange and darkgreen modules are shown. The colour of each node corresponds to module membership (MM). For each module, 20 hub genes are represented by larger-sized nodes.

To better understand these pathophysiological events, we identified genes exhibiting the most notable and consistent expression changes throughout the disease progression within each module of interest. Hub genes were defined by module membership (MM) which is a measure of the correlation between the expression pattern of a given gene and the overall expression pattern of all the genes within the module. Additionally, we derived a gene significance score from p-values obtained using DESeq2 (22)—a tool for identifying expression changes in RNA-seq data—across time stages. Strikingly, the gene significance score was found to be highly correlated (orange module R = 0.81; and darkgreen module R = 0.7) with module membership (Fig. 2B). The convergence of two different methods on the same set of genes provides evidence for the robustness of hub gene detection. The top 20 hub genes for orange and darkgreen module are labeled in Fig. 2C.

### Validation cohort confirms robustness of hub genes for muscle co-expression network in prion-infected mice

To test the reliability and validity of our findings, we investigated a validation cohort comprising samples from each time stage. We first asked whether the muscle co-expression network modules in the main cohort were preserved in the validation cohort. All modules (except for the “plum”, “orangered3”, and “brown” modules) showed high preservation with a z-summary statistic >1.96 (Supplementary Fig. 3A). We then calculated the ME for each sample in the validation cohort using the same gene module assignment as in the main cohort. Again, the “orange” and “darkgreen” modules were the most affected, exhibiting significant ME differences between RML6 and NBH at both presymptomatic and symptomatic stages (Supplementary Fig. 3B). Notably, the trend of ME changes was already evident in the early stage and consistent with the trend observed in the main cohort (Fig. 3A).

**Fig. 3.**
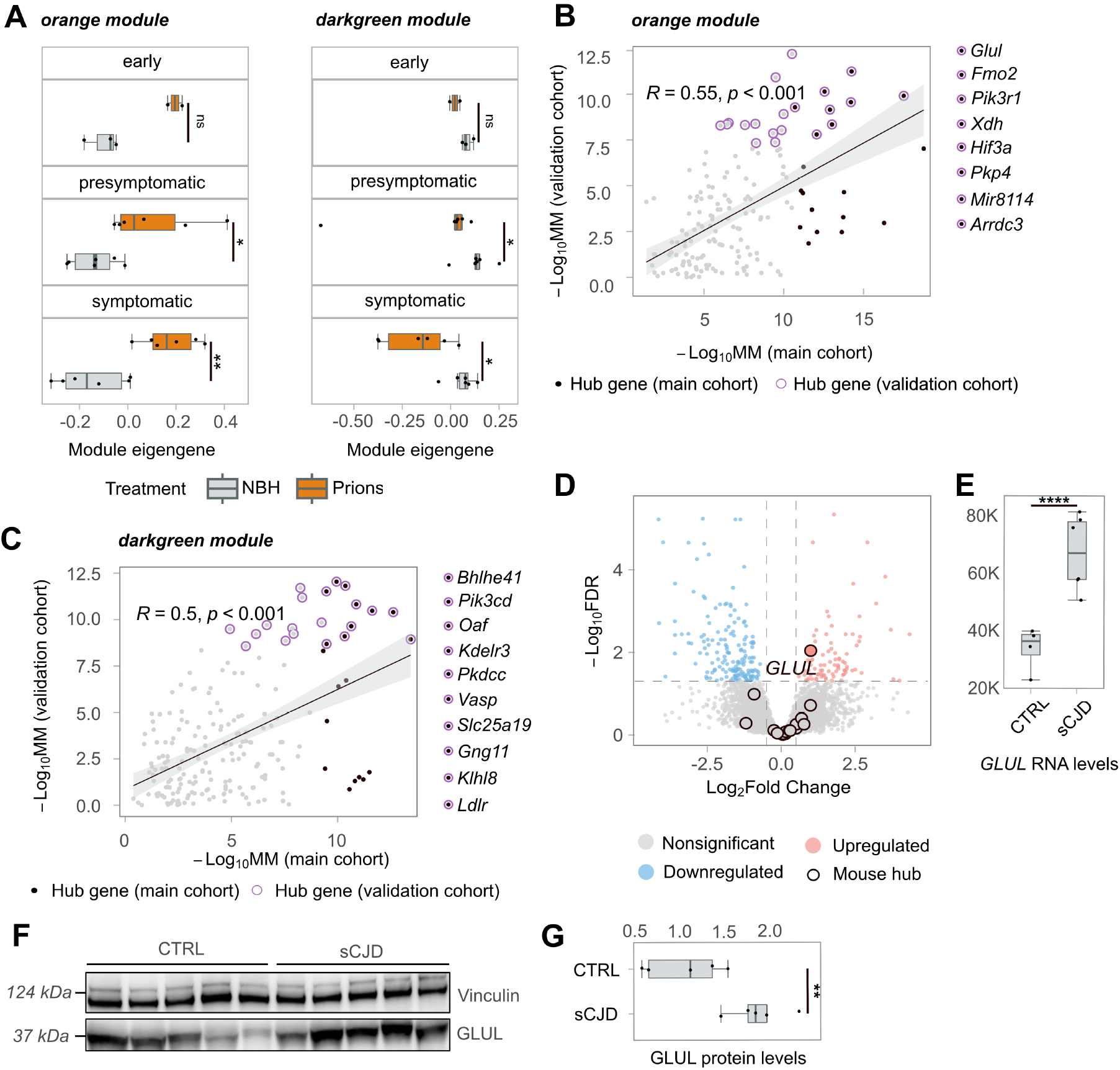
Gene Co-expression and Human Validation of GLUL Upregulation. **(A)** Boxplots of module eigengenes of the validation cohort for gene co-expression orange and darkgreen modules identified by WGCNA at different timestages (early, presymptomatic and symptomatic). **(B-C)** The scatter plots (Pearson’s correlation and pvalue indicated with R and p, respectively) depict the relationship between genes from the orange and darkgreen modules in the main and validation cohorts. Hub genes detected in the main cohort are represented by black dots, while hub genes detected in the validation cohort are represented by purple-circled dots. The black, purple-circled dots indicate hub genes detected in both cohorts. **(D)** The volcano plot displays the results of bulk RNA sequencing analysis of skeletal muscles from patients with sCJD and their age-matched controls. Red dots represent genes that are significantly upregulated in sCJD, while blue dots represent genes that are significantly downregulated. Mouse hub genes detected in the orange and darkgreen modules are black-circled. **(E)** Boxplots with normalized *GLUL* transcript counts in skeletal muscles of sCJD cases and their age-matched controls. **(F)** Western blot analysis (arbitrary densitometry unit, ADU) of GLUL and Vinculin protein expression in skeletal muscle samples from sCJD cases and age-matched controls. Each lane represents a biological replicate. **(G)** Densitometry (ADU) quantification of the Western Blot in Fig. 3F. Statistical significance (*p < 0.05, **p < 0.01, ***p < 0.005, ****p < 0.001) is indicated by asterisks.

To further validate our results, we identified hub genes for the validation cohort using MM values that correlated with those of the main cohort (orange module R = 0.55; darkgreen module R = 0.5). The overlapping hub genes between the two cohorts are shown in Fig. 3, B and C. The high correlation indicates that the hub genes identified in the main cohort were robust and reliable in the validation cohort as well. These findings reinforce the idea that the identified hub genes are biologically significant, offering insights into the underlying pathophysiological mechanisms of the disease.

### Upregulation of glutamate-ammonia ligase in skeletal muscles of prion-diseased humans and mice

To test whether our findings apply to human prion diseases, we performed RNA sequencing on skeletal muscles of sCJD patients (n = 6). For control, we used skeletal muscles of subjects without clinical or pathological diagnosis of neurodegeneration matched for age, gender, and specimen age (n = 4) (see Supplementary Table 6 for clinical details). The total RNA extracted from the Psoas major muscle exhibited significant degradation (Supplementary Fig. 4A). However, principal component analysis highlighted grouping of sCJD and control tissues in two distinct clusters (Supplementary Fig. 4, B and C). Based on |log_2_FC| > 0.5 and FDR < 0.05 (see Methods for details), we identified a total of 365 DEGs, of which 258 were protein-coding (Supplementary Table 7).

We then compared the DEGs from sCJD samples with hub genes from the two mouse cohorts, and found only one overlapping gene, glutamate ammonia-ligase (*GLUL*), which was significantly upregulated in human and mouse samples (Fig. 3D-E). We further validated the upregulation of GLUL at the protein level using Western Blot (Fig. 3F-G).

While examining post-mortem muscle samples from human patients with sporadic Creutzfeldt-Jakob disease (sCJD), we found numerous changes in gene expression that differed from those observed in mouse models. These differences might have only become apparent during the late stages of the disease, where nonspecific changes, possibly influenced by factors like prolonged immobilization, are more likely to occur. However, our analysis of mouse models provides strong evidence that GLUL correlates with the disease process at earlier stages.

### GLUL is upregulated in mice infected with a variety of prion strains

Prion strains are infectious isolates exhibiting distinct biological properties, such as tissue tropism, incubation time, and neuropathological features (23). Different prion strains may elicit different transcriptional responses in their hosts. To determine whether *Glul* alterations are a common feature across prion strains, we tested whether *Glul* is changed in mice infected with a panel of mouse-adapted prion strains. To this end, we intracerebrally injected an additional cohort of C57BL/6 mice with ME7, 22L, RML6, and NBH, and collected hindlimb skeletal muscle at different time points (8 wpi, 16 wpi and terminal) corresponding to each of the three aforementioned disease stages.

At 8 wpi, *Glul* RNA and protein levels were not altered in both ME7 and 22L strains. At the protein level, the most substantial upregulation was found in RML6-infected animals (Fig. 4, A-C). The delayed *Glul* upregulation in mice inoculated with the ME7 and 22L strains may be related to differences in disease onset, as RML6 induced disease more quickly than ME7 and 22L. *Glul* expression was significantly altered at both the RNA (Supplementary Table 7) and protein level in RML6 and ME7 strains, while only at RNA level in 22L strain at 16 wpi. In terminally sick mice, both RNA and protein level of Glul were upregulated in all strains (Fig. 4, A-C). These results suggest that Glul upregulation is a universal feature across various prion diseases, highlighting its potential role in the underlying pathophysiological processes of these conditions. Such a consistent pattern of dysregulation suggests that glutamate ammonia ligase may influence disease progression and clinical manifestations in prion-affected individuals.

**Fig. 4.**
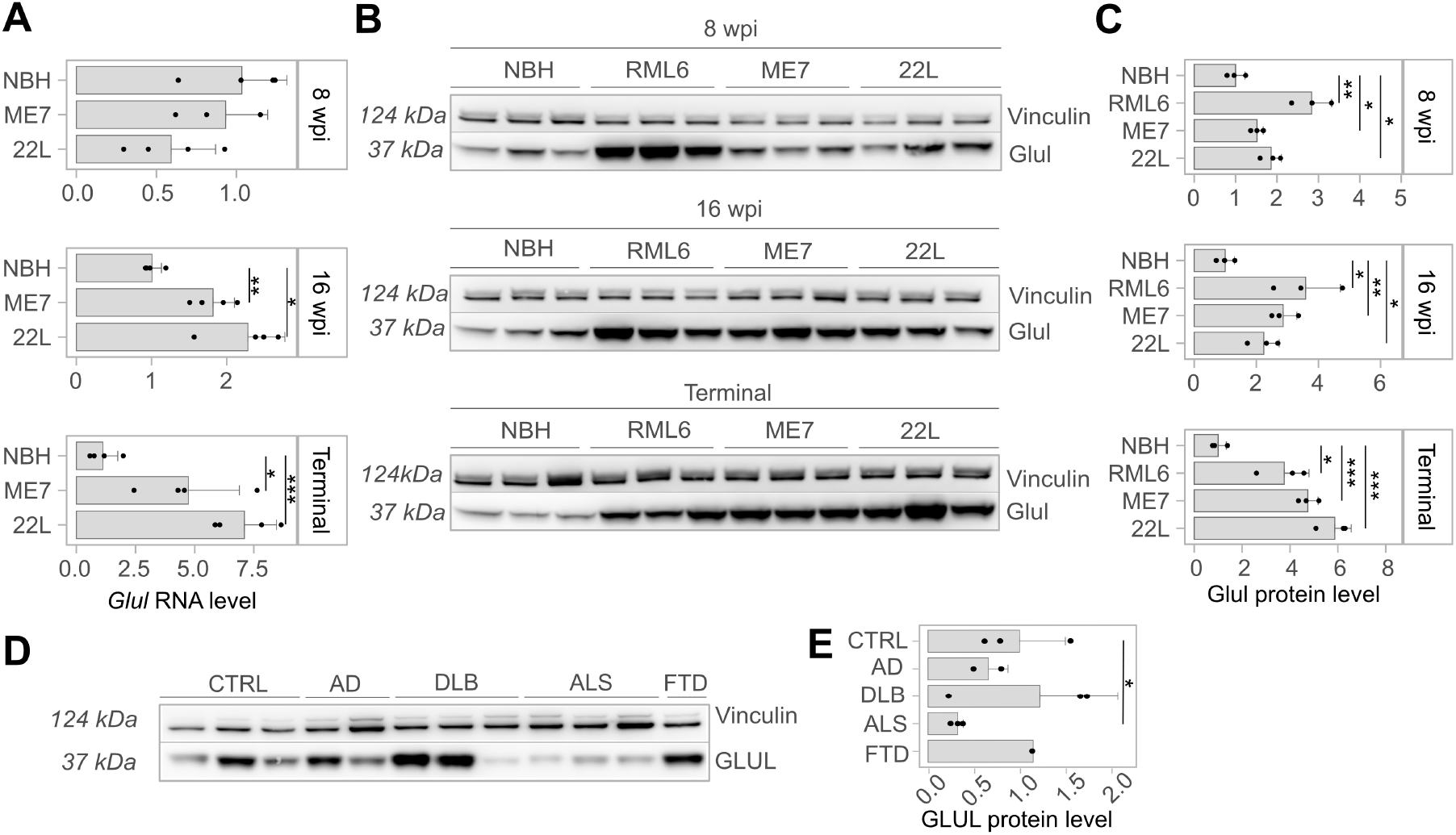
Levels of Glul mRNA, protein, glutamate, and glutamine in skeletal muscle lysates at 8 and 16 weeks post-inoculation (wpi) and terminal stage of mice with prion strains RML6, ME7, and 22L, as well as related control (NBH). In panel **(A)**, barplots display *Glul* mRNA levels normalized by GAPDH mRNA levels (derived from Ct values via RT-PCR). **(B)** Western blots of Glul and Vinculin protein levels of infected with different prion strains, as well as related NBH control **(C)** Densitometry (arbitrary densitometry unit, ADU) quantification of the Western Blot in Fig. 4B. **(D)** Western blot of Glul and Vinculin protein levels of skeletal muscles from AD, DLB, ALS and FTD diagnosed individuals **(E)** Densitometry (ADU) quantification of the Western Blot in Fig. 4D. Statistical significance (*p < 0.05, **p < 0.01, ***p < 0.005, ****p < 0.001) is indicated by asterisks.

### GLUL upregulation is specific to prion diseases

To further investigate the disease specificity of *Glul* dysregulation, we examined its protein levels in hindlimb skeletal muscles of mouse models for ALS, AD, and DLB. Interestingly, Glul did not exhibit upregulation in these models. The confinement of Glul upregulation to prion diseases points to specific pathogenetic mechanisms that do not occur in other types of neurodegenerations (Supplementary Fig. 5C, D). To corroborate these findings, we extended our investigation to skeletal muscle necropsies from patients with familial ALS, FTD (Frontotemporal Dementia), AD, and DLB. Again, GLUL protein levels were unaltered in AD, DLB and FTD, highlighting its distinct role in prion diseases compared to other neurodegenerative conditions. Notably, GLUL protein levels were significantly downregulated in all three individuals affected by ALS (Fig. 4D-E).

### Glutamate-glutamine biosynthesis in muscle is affected during prion disease progression

Glul catalyzes the conversion of glutamate to glutamine, whereas glutaminase carries out the opposite process. Given the increased *Glul* expression in skeletal muscle during prion disease progression, we investigated whether this metabolic pathway was affected. We measured glutaminase, glutamate and glutamine levels in muscle lysates from C57BL/6 mice inoculated with ME7, 22L and RML6 brain homogenates, as well as NBH for control. At the early disease stage, there were no significant alterations in the levels of glutamate and glutamine (Supplementary Fig. 6A, E), even in RML6-infected animals, despite the increased expression levels of Glul. This might be attributable to a compensatory effect of glutaminase upregulation (Supplementary Fig. 5A-B) which may maintain glutamate homeostasis and offset the increased expression of Glul. Notably, we observed an increase in the expression of Glul at 16 wpi (two weeks prior to the onset of clinical signs of prion disease). Despite the Glul upregulation, the balance of glutamate and glutamine was maintained (Supplementary Fig. 6A; E), as well as glutaminase levels (Supplementary Fig. 5A, B), pointing to a metabolic ability restoring the levels of glutamate consumed by Glul upregulation. At the terminal stage of the disease, the levels of glutamate were conspicuously reduced (Supplementary Fig. 6A), plausibly due to a significant upregulation of Glul. However, the levels of glutamine and glutaminase were unchanged (Supplementary Fig. 6E; Supplementary Fig. 5A). Glutamate reduction was also detected in muscle necropsies of sCJD patients versus age- and sex-matched controls (Supplementary Fig. 6B) despite unchanged glutamine levels (Supplementary Fig. 6F).

We then examined the levels of glutaminase, glutamate, and glutamine in skeletal muscles of aged mouse models for AD, DLB, and ALS. Unlike in prion diseases, we found no significant alterations in glutaminase (Supplementary Fig. 5C, E), glutamate (Supplementary Fig. 6C), and glutamine (Supplementary Fig. 6G) levels in these neurodegenerative disorders. This highlights the specificity of the observed metabolic changes in prion diseases. We then evaluated glutamate and glutamine levels in skeletal muscles of human AD, DLB, ALS and FTD cases. Our results revealed a reduction in glutamate levels solely in the context of ALS, mirroring the observed trend in prion diseases (Supplementary Fig. 6, D and H). However, it is important to note that while GLUL expression was downregulated in ALS, it was upregulated in prion diseases.

## Discussion

Transcriptional changes observed in the early phases of prion disease offer valuable insights into the pathogenic mechanisms at play in extraneural organs. These alterations in gene expression not only enhance our understanding of the disease’s progression but also underscore the systemic nature of prion diseases (PrDs), revealing critical aspects of their pathology beyond the central nervous system. Here we conducted a comprehensive transcriptomic characterization of extraneural organs known to harbor prions. We hypothesized that the presence of prions in these organs could result in changes to RNA processing, or modifications to the abundance of specific transcripts. By examining multiple timestages throughout the progression of the disease in prion-inoculated mice, this investigation provides insights into the pathological processes preceding the onset of clinical symptoms. The selection of timestages was based on a recent study where authors defined disease stages based on clinically relevant EEG (electroencephalography): recordings between NBH and RML mice groups began to diverge at 10 wpi and became significantly different at 18 wpi marking the beginning of clinical onset (21). Other authors defined symptomatic stage as the period from 18 wpi to terminal stage based on motor function impairments (20).

Significant changes in gene expression were observed in all three organs. In the spleen, major transcriptional alterations were detected only at the terminal stage of the disease, which suggests a slower accumulation rate of prions in lymphatic tissue following intracerebral inoculation compared to ingestion or intraperitoneal inoculation. Conversely, blood samples displayed upregulation of numerous transcripts during the early stages of prion exposure, of which a significant fraction was related to hemostasis.

Gene-expression changes were detected in both blood and spleen but were not consistent across all timepoints. However, the transcriptome of skeletal muscle showed consistent alterations throughout the disease progression. Using WGCNA, we identified two primary gene subsets in skeletal muscle exhibiting progressive changes that became evident in the early stage of disease. Validation of these findings in an independent cohort of mice, in which sequencing libraries were prepared several months after RNA extraction from snap-frozen skeletal muscles by a different researcher, enhances the robustness of the results. This validation underscores the consistency of the findings even in the face of significant technical variability introduced by the procedures. These results have potential applications in monitoring disease progression following therapeutics administration, in conjunction with molecular and behavioral assessments for evaluating treatment efficacy (24–26).

We evaluated the relevance of our findings in humans by conducting RNA sequencing on skeletal muscle samples obtained from individuals with sCJD and control individuals without neurological impairments. While we were unable to confidently evaluate the level of preservation of mouse and human muscle gene-expression networks, we found little overlap between the joint set of hub genes identified across two mouse cohorts and human DEGs. There may be various reasons for this discrepancy, including genetic differences between mice and humans resulting in varying biological responses to similar diseases (27). Additionally, different conditions were employed to process the human samples and decontaminate them from prions, which may have influenced the results (28). Variations in disease stage or severity, as well as differences in the tissue types analyzed (hindlimb muscle in animals vs. psoas muscle in humans), could have also played a role in the observed differences (29). Finally, there may be underlying differences in the disease biology between humans and mice that are not yet fully understood or characterized (30). Despite the abovementioned limitations in translating mouse findings to human ones, we found that *GLUL*, which was a strong hub gene in the upregulated “orange” module in the mouse study, was significantly upregulated in human sCJD samples, as confirmed at the protein level by Western Blot analysis. Hence, the consistent upregulation of GLUL may be indicative of its significant role in the pathophysiology of human prion diseases, potentially contributing to the progression of the disease, especially in its early stages. GLUL is a glutamate-ammonia ligase that catalyzes the synthesis of glutamine from glutamate and ammonia in an ATP-dependent reaction.

There is some evidence that *GLUL* expression in the brain is not altered in PrDs. *Glul* mRNA levels were unchanged in the brains of mice infected with prions compared to uninfected mice (20). Another recent study found that *GLUL* expression was also unchanged in the brains of patients with sCJD compared to healthy controls (31). Therefore, the alterations in glutamate/glutamine metabolism in prion-infected brains (32) that were proposed to contribute to neurodegeneration and cognitive dysfunctions, are unlikely to stem from any changes in brain-resident GLUL. Instead, *Glul* expression was consistently upregulated in the skeletal muscles of animals throughout the disease, and this finding was present after infection with multiple prion strains. This suggests that *GLUL* may also be upregulated before the clinical onset of human PrDs. The identification of a common pathological phenotype among these diseases would be a significant finding, shedding light on the underlying mechanisms specific to these conditions at early stages. It is important to note that this phenomenon appears to be unique to these diseases and does not occur in other common neurodegenerative disorders such as ALS, AD, or DLB; interestingly, GLUL protein levels were consistently found to be downregulated in human ALS. This distinct pattern underscores the unique role of GLUL in the pathophysiology of prion diseases, differentiating them from other neurodegenerative conditions in terms of their molecular and cellular mechanisms.

We hypothesized that the upregulation of GLUL protein in skeletal muscle may be linked to a metabolic dysfunction in the glutamate-glutamine pathway. Prior research had shown the direct influence of glutamine levels on the expression of *GLUL* in skeletal muscle cells (33). Building upon this finding, it is plausible to speculate that a potential systemic deficit in glutamine levels might contribute to the observed upregulation of *GLUL* in skeletal muscle. Skeletal muscle exerts a pivotal role in glutamine storage, production, and release into the bloodstream. This hypothesis gains further support from the significant reduction in glutamate levels observed, which could be attributed to the heightened activity of GLUL in response to altered glutamine availability. This was unexpected as previous studies had reported increased levels of glutamate and glutamine in the brain and cerebrospinal fluid of patients with neurodegenerative diseases, including prion diseases (32). The regulation of glutamate and glutamine metabolism in skeletal muscle differs from that in the brain, and the upregulation of *GLUL* in skeletal muscle may have a distinct role in the pathology of prion disease. In contrast, the absence of such *GLUL* upregulation in other NDDs might signify a distinct metabolic response. The lack of compensatory *GLUL* expression could contribute to the sustained alterations in glutamine and glutamate levels seen in those conditions. Therefore, the specific association between *GLUL* upregulation and prion diseases suggests a unique interplay between glutamine/glutamate metabolism and disease progression, setting prion diseases apart from other NDDs. Finally, extraneural GLUL measurements may represent an accessible biomarker for monitoring the efficacy of experimental antiprion therapies.

## Materials and Methods

### Animals

Prion-inoculated and control-injected mice were regularly monitored for the development of clinical signs, according to well-established procedures using humane termination criteria. Intracerebral injections and transcardiac perfusions were performed in deeply anesthetized mice. Habituation periods before the experiment began were included. Male C57BL/6J mice were obtained from Charles River, Germany. Mice were housed in a conventional sanitary facility and monitored for the presence of all viral, parasitic, and bacterial species listed in the Federation of European Laboratory Animal Associations (FELASA). The facility was tested positive for Murine Norovirus and Helicobacter spp. The mice were housed in IVC type II long cages and up to five animals were housed in the same cage which were staffed with individual apartments. Mice had unrestricted access to sterilized drinking water and were fed *ad libitum* a pelleted mouse diet. The light/dark cycle consisted of 12/12 h with artificial light. The temperature in the room was 21 ± 1 °C with a relative humidity of 50 ± 5%. Air pressure was regulated at 50 Pa, with 15 complete changes of filtered air per hour by a HEPA filter.

### Prion inoculations and processing of tissue samples

Animals, prion inoculation and necropsy procedures are identical to those described in (20). C57BL/6J male mice were inoculated in the right hemisphere with either 30 µl of passage 6 of Rocky Mountain Laboratory (RML6), or 22L, or ME7 strain mouse-adapted scrapie prions containing 9.02 LD_50_ of infectious units per ml in 10% w/v homogenate. Non-infectious brain homogenate (NBH) from CD1 mice was used as a negative control. Mice were assigned randomly to experimental groups. Animals were monitored at least thrice per week, after the clinical onset of prion disease, they were monitored daily, and prion-inoculated mice were terminated upon evident signs of terminal disease. NBH-inoculated mice were sacrificed 13 days after the termination of the last prion-inoculated mice. Whole blood, spleen and muscle were dissected, snap-frozen in liquid N_2_ and stored at −80 °C prior to sequence library generation.

### Processing of AD, DLB and ALS tissue samples

Double-transgenic APP/PS1 mice (n = 3; 8 months of age) were used as AD mouse models from which we have collected hindlimb skeletal muscles. For the DLB mouse model, hindlimb skeletal muscles of transgenic A53T synuclein mutant mice lines were kindly provided by Dr. Noain Daniela’s group (Department of Neurology, University Hospital Zurich; n=2, 8 months of age) and by Dr. Ruiqing Ni’s group (Institute for Biomedical Engineering, University Hospital Zurich; n=1, 8 months of age). Non-transgenic C57BL/6J male littermates (n = 3, 8 month old) were used as controls. All mice were housed under a 12-hour light/12-hour dark schedule and had free access to food and water. All animals were euthanized by pentobarbital injection. Skeletal muscles were dissected, snap frozen in liquid N_2_ and stored at −80 °C prior to western blot and biochemical analyses.

Differently, skeletal muscle lysates obtained from both wild-type and SOD1^G93A^ transgenic mouse models of ALS were generously provided by Prof. Dr. Musarò, the Principal Investigator leading the neuromuscular research group at the Sapienza University of Rome. To provide a concise overview, wild-type C57BL/6 (WT) and transgenic SOD1^G93A^ mice were utilized for our investigation. The mice were sacrificed at 130–140 days of age, a time point closely aligned with the spontaneous mortality of SOD1^G93A^ mice. Euthanasia was conducted through cervical dislocation to ensure minimal suffering. Immediately following the humane sacrifice, muscle samples were excised for subsequent analysis, with one muscle specimen collected from each animal for testing purposes. Tissue lysates were prepared according to (34).

### Preparation of RNA libraries for Mouse sequencing

RNA was extracted from snap-frozen organs by means of the RNeasy Plus Universal Kit (QIAGEN). The quantity and quality of RNA were analyzed with Qubit 1.0 Fluorometer (Life Technologies) and Bioanalyzer 2100 (Agilent Technologies), respectively. For library preparation, we used TruSeq RNA Sample Prep kit v2 (Illumina). We performed poly-A enrichment on 1 μg of total RNA per sample, which was then reverse transcribed into double-stranded cDNA followed by ligation of TruSeq adapters. Sequencing fragments containing TruSeq adapters at both termini were enriched by PCR. Quantity and quality of enriched libraries were analyzed using Qubit (1.0) Fluorometer and Caliper GX LabChip GX (Caliper Life Sciences), which showed a smear corresponding to a mean fragment size of around 260 bp. Libraries were then normalized to 10 nM in Tris-Cl 10 mM, pH 8.5, with 0.1% (v/v) Tween 20. Cluster generation was performed with the TruSeq PE Cluster kit v4-cBot-HS (Illumina), using 2 pM of pooled normalized libraries on the cBOT. Sequencing was performed on Illumina HiSeq 4000 paired-end at 2 × 126 bp using the TruSeq SBS kit v4-HS (Illumina).

### Patient samples

Human skeletal muscle samples of psoas major muscle were collected from patients with a clinical suspicion of Creutzfeldt-Jakob’s disease and submitted for an autopsy to the Swiss National Reference Center for Prion Disease between 2004 - 2011. A detailed description of samples used for RNA extraction and sequencing is given in Supplementary Table 8. Sporadic CJD was diagnosed according to criteria described previously (35).

### Preparation of RNA libraries for Human sequencing

Firstly, CJD bulk tissues were lysed in TE buffer with the anionic detergent sodium dodecyl sulphate (SDS) and digested at 50°C with 2 mg / ml^-1^ Proteinase K (PK) for 2 hours to eliminate solids and release DNA/RNA from proteins. Although prions are well-known for their relative resistance to PK digestion, prion infectivity largely depends on PK-sensitive oligomers. Indeed, prolonged PK digestion reduces prion titers by a factor of >10^6^, but residual PK-resistant material may still be infectious. In a second step, TRIzol reagent solution was added to the lysate (it contains Gdn-SCN and phenol, which inactivate RNases and disaggregate prions) and kept overnight at 4°C. Gdn-SCN is a chaotropic salt which rapidly denatures proteins and abolishes the infectivity of prion disease inoculum. At high concentrations, guanidine salts disaggregate PK-resistant PrP^Sc^ fibrils, eliminate PK resistance and abolish PrP^Sc^ conversion, meaning that any PK-resistant material that survived the digestion step would be expected to be inactivated at this stage of the protocol. 0.2 ml of ultrapure phenol:chloroform:isoamyl alcohol (Thermo Fischer Scientific) was added to the samples, followed by strong shaking and incubation at room temperature for 5 mins. Centrifugation step at 12,000 x g for 15 min at 4 °C generated two phases. The aqueous upper phase was transferred to a fresh tube; 0.5 ml of isopropanol and 1 µl of Glycoblue Coprecipitant (Thermo Fisher Scientific) were added. Next, RNA was pelleted for 20 min at 12,000 x g at 4°C and washed twice with 75% ethanol. The RNA pellet was dissolved at 55 °C in 20 µl of free nuclease water.

The quantity and quality of RNA were analyzed with Qubit 1.0 Fluorometer (Life Technologies) and Tapestation 4200 (Agilent Technologies), respectively. The TruSeq stranded RNA protocol (Illumina) was employed for library preparation. In brief, 1 μg of total RNA per sample was poly-A enriched, reverse transcribed into double-stranded cDNA and then ligated with TruSeq adapters. PCR was performed to selectively enrich for fragments containing TruSeq adapters at both ends. The quantity and quality of enriched libraries were analyzed using Qubit (1.0) Fluorometer and Tapestation 4200. The resulting product is a smear with a mean fragment size of approximately 260 bp. Libraries were then normalized to 10 nM in Tris-Cl 10 mM, pH 8.5, with 0.1% (vol/vol) Tween 20. Cluster generation was performed with the TruSeq PE Cluster kit v4-cBot-HS (Illumina), using 2 pM of pooled normalized libraries on the cBOT. Sequencing was performed on Illumina HiSeq 4000 paired-end at 2 × 126 bp using the TruSeq SBS kit v4-HS (Illumina).

Despite profound RNA degradation, we speculated that moderate RNA degradation might preserve biological information. We then decided to set a RIN ≥ 3 as a minimal threshold for human muscle tissue. 12 out of 28 initially collected samples passed the RIN threshold (Supplementary Fig. 4A and Supplementary Table 4) and were further processed. 2 out of 12 samples were subsequently removed because of a high cluster condition variance (upon the quality control of the sequencing) resulting in a final sample size of n = 10 for downstream analysis.

### Differential gene expression

We used FASTQC and parallel (36) algorithms for quality control of raw sequencing reads. We clipped low-quality ends as follows: 5’ = 3 bases, 3’ = 10 bases. Reads were aligned to mouse mm10 and human GRCh38.p13 reference genome, and transcriptome using STAR v2.3.0e_r291 (37) on cloud computing solution SUSHI of the Functional Genomics Center of Zurich (38). DESeq2 (22) was used to detect differentially expressed genes based on the following thresholds: (a) |log_2_-fold change| > 0.5 (b) FDR < 0.05. Genes with less than 10 counts in total were excluded. Sex was included as a covariate in the formula for analyzing human samples with DESeq2. Gene ontology analysis was performed using clusterProfiler for R (39).

### WGCNA

WGCNA was performed using the WGCNA R package (40). Outlier genes were identified and removed using the *goodSamplesGenes()* function. Additionally, genes with fewer than 10 counts in over 50% of samples were filtered out. Raw count data was normalized using the variance stabilizing transformation provided by the DESeq2 R package. An adjacency matrix was generated using the *adjacency()* function with default parameters. To meet the criteria for a scale-free network, a soft threshold of 4 was uniformly applied to all networks. Adjacency matrix was transformed into a Topological Overlap Matrix (TOM). Average linkage hierarchical clustering was performed on a dissimilarity matrix (1 – TOM) and subsequently, modules of co-expressed genes were identified using the dynamic cut tree algorithm (*cuttreeDynamic function()*), with a minimum cluster size set to 30. Similar modules were merged based on their module eigengene (ME) correlation. To assess the significance of differences in ME values between tested conditions, a Mann-Whitney U test was conducted. To identify genes with the highest connectivity within modules, Module Membership (MM) was computed as the Pearson correlation coefficient (p-value) between individual gene expression levels and the ME. For genes in the modules of interest, we calculated a gene significance score based on the p-values calculated with DESeq2 for each timestage. Specifically we combined p-values for different time stages using the *combineParallelPValues()* function from the metapod R package with the method argument set to “stouffer” (41). Negative log base 10 of the combined p-value represents the gene significance score. The preservation of mouse muscle co-expression network was tested using *modulePreservation()* function from the WGCNA package, using the network from the main cohort as a reference.

### Western blot analysis

To prepare the samples, 1 ml of cell-lysis buffer (20 mM Hepes-KOH, pH 7.4, 150 mM KCl, 5 mM MgCl2, 1% IGEPAL) supplemented with protease inhibitor cocktail (Roche 11873580001) was added to the lysed samples. They were then homogenized twice at 5’000 rpm for 15 seconds using a Precellys24 Sample Homogenizer (LABGENE Scientific SA, BER300P24) and incubated on ice for 20 minutes. The cleared lysates were obtained by centrifugation at 2’000 rcf, 4° C for 10 minutes in an Eppendorf 5417 R tabletop centrifuge. The concentration of whole protein was determined using a BCA assay (Thermo Scientific). The samples were boiled in 4 x LDS (Invitrogen) containing 10 mM DTT at 95°C for 5 minutes. 15 µg of total protein per sample were loaded onto a 4–12% Novex Bis-Tris Gel (Invitrogen) gradient for electrophoresis at 80 V for 15 minutes, followed by constant voltage of 150 V. The PVDF or Nitrocellulose membranes were blocked with 5% Sureblock (LubioScience) in PBS-T (PBS + 0.2% Tween-20) for 1 hour at room temperature. Membranes were then cut into three parts according to the molecular weight. The membrane was divided into three segments for targeted antibody incubation: The upper portion was treated with anti-Vinculin (1:5000, Abcam, ab129002), the middle section with anti-Glutaminase (1:3000, Abcam, ab202027), and the lower segment with anti-Glutamine synthetase (1:2000, Abcam, ab176562). This antibody incubation was carried out in PBS-T supplemented with 1% Sureblock, and the membranes were left overnight at 4°C to facilitate optimal binding. They were washed thrice with PBS-T for 10 minutes. The membranes were incubated with secondary antibodies conjugated to horseradish peroxidase (HRP-tagged goat anti-rabbit IgG (H+L), 1:3000, 111.035.045, Jackson ImmunoResearch) for 1 hour at room temperature. The membranes were washed thrice with PBS-T for 10 minutes and developed using a Classico chemiluminescence substrate system (Millipore). The signal was detected using a LAS-3000 Luminescent Image Analyzer (Fujifilm) and analyzed with ImageJ software.

### Biochemical analysis

The concentration of glutamine and glutamate in skeletal muscle was measured using the Merck Glutamine Assay Kit (Catalog Number MAK438). To prepare the lysates, a total of 600 µg of total protein was utilized. In each standard and sample well, 80 µL of working reagent was added to determine the glutamine concentration. To measure glutamate concentration, samples were also prepared with 80 µL of blank working reagent. The 96-well plate was incubated at 37°C for 40 minutes. Absorbance values were recorded at 450 nm using a microplate reader. The concentrations of glutamine and glutamate were determined by comparing the absorbance of the samples to a standard curve generated from known concentrations of glutamine and glutamate. The results were expressed as µM/ml.

### Study approval

Animal experiments were approved by the Veterinary Office of the Canton Zurich (permit numbers ZH41/2012, ZH90/2013, ZH040/15, ZH243/15) and carried out in compliance with the Swiss Animal Protection Law. Animal discomfort and suffering was minimized as much as possible and individual housing was avoided.

We obtained sCJD anonymized skeletal muscle samples from an approved study sanctioned by the Cantonal Ethics Committee of the Canton of Zurich under approval number #2019-01479 and a written informed consent was received prior to participation.

The experimental protocols involving GLUL’s specificity validation test on other neurodegenerative diseases by using human participants adhered strictly to the guidelines set by French regulations. Prior to participation, comprehensive written informed consent was diligently obtained from all individuals, including those who underwent skeletal muscle necropsies for Familial Amyotrophic Lateral Sclerosis (Familial SLA), Fronto-Temporal Dementia with ALS (DFT-SLA), and pure cases of Alzheimer’s Disease (AD) and Dementia with Lewy Body (LBD). Human biological samples and associated data were obtained from Tissu-Tumorothèque Est (CRB-HCL, Hospices Civils de Lyon Biobank, BB-0033-00046).

## Data availability

Raw sequencing data as well as processed data from this manuscript is available freely via GEO accession number GSE210128. Code to reproduce results generated in this manuscript will be available upon publication at https://github.com/marusakod/RML_extraneural_organs.

## Author’s contributions

DC performed CJD’s RNA extraction, sample preparation and analyses, performed all the experimental validations on both animal models and human samples and wrote and reviewed the manuscript. MK analyzed the sequencing data, conceptualized, and developed the shiny app, wrote, and critically reviewed the manuscript. KF analyzed and visualized initial sequencing data, acquired funding, and critically reviewed the manuscript. SS and MN conceived the study. SS, MN, and JB performed an initial analysis of the data, extracted RNA, prepared the sequencing library in cooperation with the Functional Genomics Center Zurich of the ETH Zurich and the University of Zurich, supervised personnel, and critically reviewed the original manuscript. PS bred the animals, performed inoculations, clinical scoring, harvested organs and was responsible for other important tasks related to animal husbandry. SS performed human-related samples validation and critically reviewed the original manuscript. NS made the final diagnosis of DLB, AD and ALS individuals, and performed skeletal muscle biopsies. CS conceived the study, performed initial data analysis, supervised personnel, and reviewed the original manuscript. AA conceived the study, provisioned animals, personnel and IT resources, supervised personnel, provided advice on experiments, coordinated the project, acquired funds, wrote parts of the manuscript, and critically reviewed the final manuscript version.

## Acknowledgements

KF is a Visiting Faculty Program Fellow at the Department of Molecular Neuroscience, Weizmann Institute of Science, hosted by Prof. Eran Hornstein, and receives unrestricted funding from the Benoziyo Endowment Fund for the Advancement of Science and Swiss Society of Friends of the Weizmann Institute of Science. AA is supported by an Advanced Grant of the European Research Council (ERC, No. 250356), a Distinguished Scientist Award of the Nomis Foundation, grants from the GELU foundation, the Swiss National Foundation (SNF, including a Sinergia grant, #179040, 183563, 207872), and the Swiss Initiative in Systems Biology, SystemsX.ch (PrionX, Synucleix). The authors are grateful to Martina Cerisoli, Nicola Conneely, Andrea Armani, Marigona Imeri, and Mirzet Delic for support, assistance in laboratory investigations and animal husbandry, and also Med. Dr. Regina Reimann for providing sCJD skeletal muscles biopsies. The authors acknowledge the Functional Genomics Center Zurich of the ETH Zurich, and Next Generation Sequencing Platform of University of Bern for preparing sequencing libraries, RNA sequencing, quality control and technical support of mouse and human studies, Prof. Eli Eisenberg (Raymond and Beverly Sackler School of Physics and Astronomy and Sagol School of Neuroscience, Tel Aviv University, Tel Aviv, Israel) for help with RNA editing analyses. The authors extend their gratitude to Prof. Musarò Dr. Gabriella Dorbowolny and Dr. Gaia Laurenzi (La Sapienza University), for providing SOD1^G93A^ muscle lysates. Appreciation is also extended to Dr. Daniela Noain, Ines Antunes dos Santos Dias and Irena Barbaric (Department of Neurology, University of Zurich) for their kind contribution of DLB hindlimb skeletal muscles and Dr. Ruiqing Ni’s group and Benjamin Francois Combes (Institute for Biomedical Engineering, University of Zurich) for providing DLB hindlimb skeletal muscles. We acknowledge the Tissu-Tumorotheque Est (CRB HCL, HCL’s biobank) for providing the human biological samples (AD, DLB, ALS, FTD) used in this study. The funders of the study had no role in study design, data collection, data analysis, data interpretation, or writing of the report. The corresponding author had full access to all the data in the study and had final responsibility for the decision to submit for publication.

---

**Supplementary Fig. 1.**
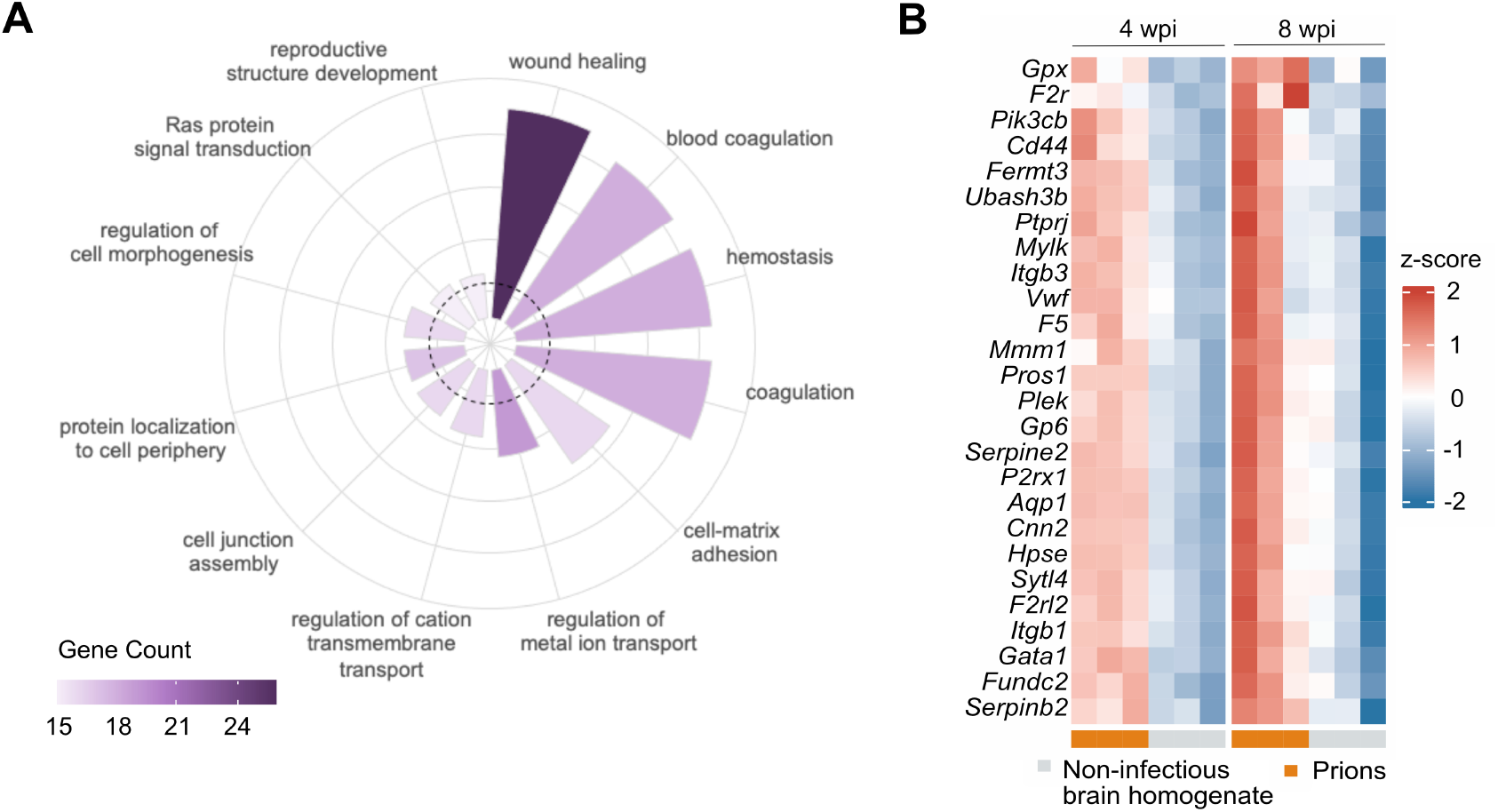
Early Changes in blood associated with hemostasis and wound healing terms. **(A)** The 270 overlapped, upregulated, blood-derived DEGs at 4wpi and 8wpi are associated with specific Gene Ontology (GO) terms. The Randarplot displays the results of the GO over-representation analysis by Biological Process (BP) ontology class. **(B)** Expression patterns (z-score based) of genes related to hemostasis process in blood at 4 and 8 wpi.

**Supplementary Fig. 2.**
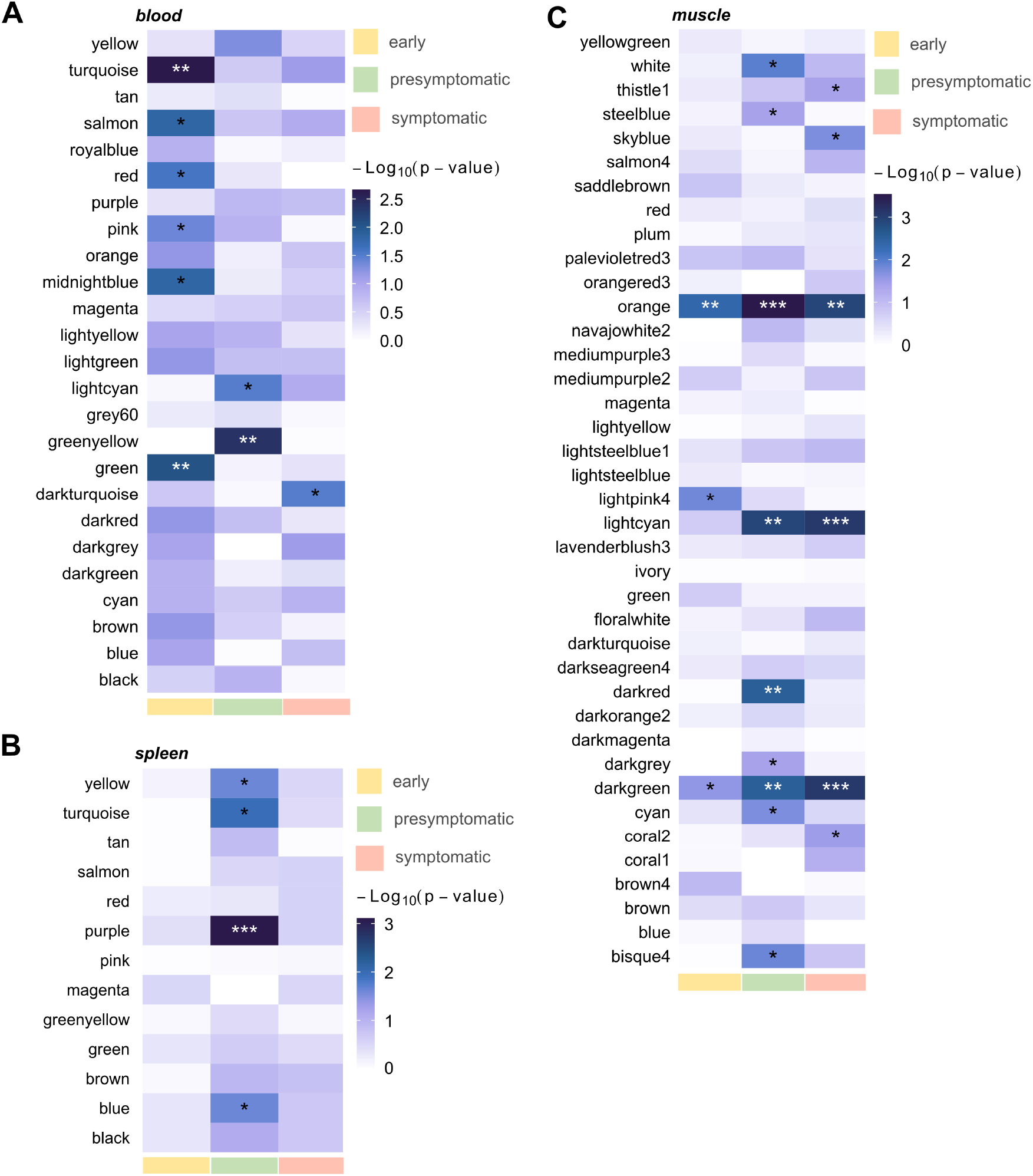
Modul*e* Eigengene Significance Heatmap for Different Organs and Time Stages. Each heatmap illustrates module eigengene significance at three timestages (early, pre-symptomatic, and symptomatic) derived from the comparison between NBH and RML6 inoculated mice from **(A)** whole blood, **(B)** spleen, and **(C)** skeletal muscles necropsies. Each row represents a specific module, while columns correspond to individual timestages. Statistical significance (*p < 0.05, **p < 0.01, ***p < 0.005, ****p < 0.001) is indicated by asterisks.

**Supplementary Fig. 3.**
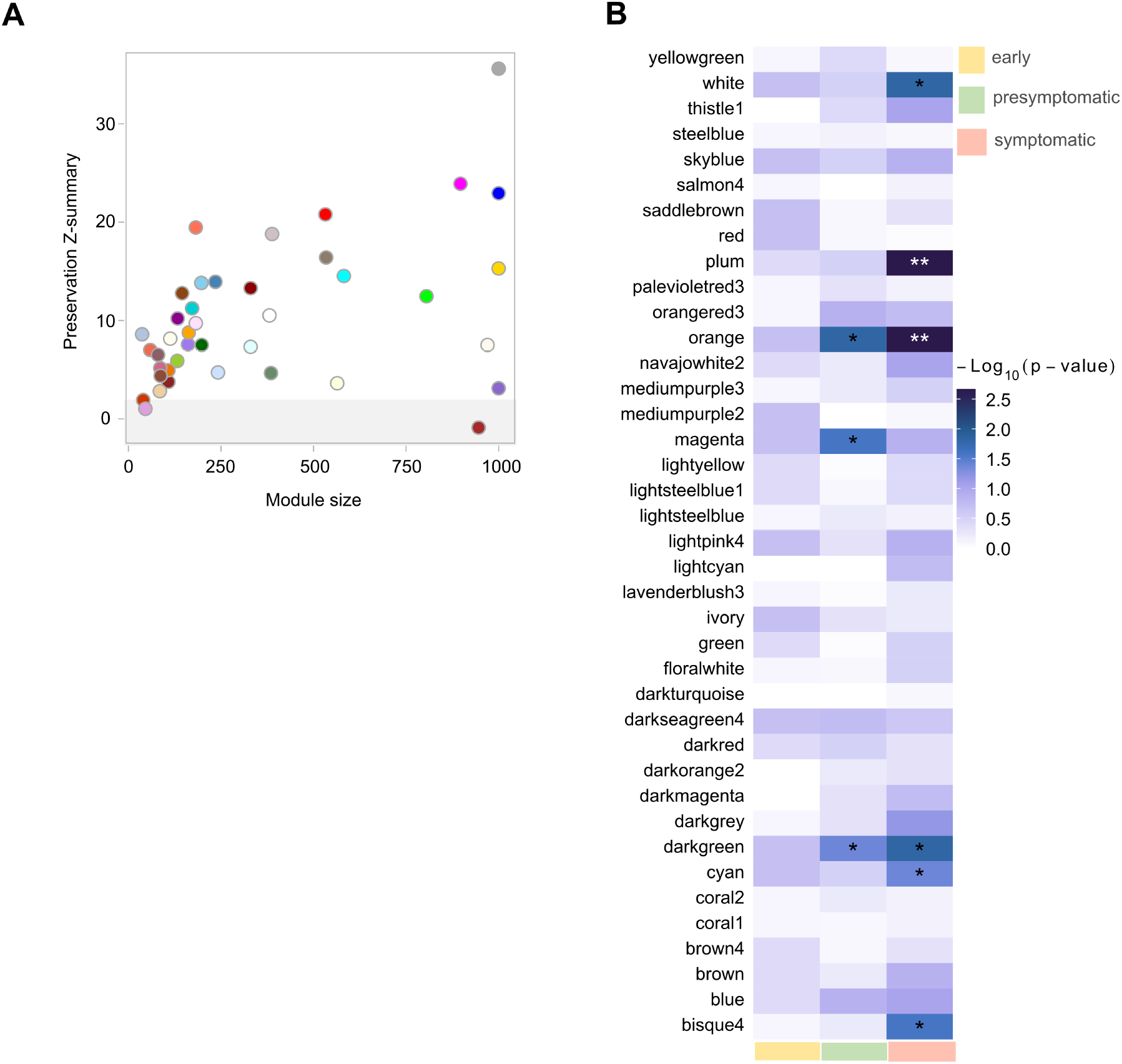
Module Preservation, and Module Eigengene of validation cohort. **(A)** Scatter plot of the Z-summary module preservation statistic and the sizes of modules in muscle co-expression network. The modules with Z-summary > 1.96 were interpreted as preserved. **(B)** The heatmap illustrates module eigengene significance at three timestages (early, pre-symptomatic and symptomatic) derived from the comparison between NBH and RML6 inoculated mice. Each row represents a specific module, while columns correspond to individual timestages. Statistical significance (*p < 0.05, **p < 0.01, ***p < 0.005, ****p < 0.001) is indicated by asterisks.

**Supplementary Fig. 4.**
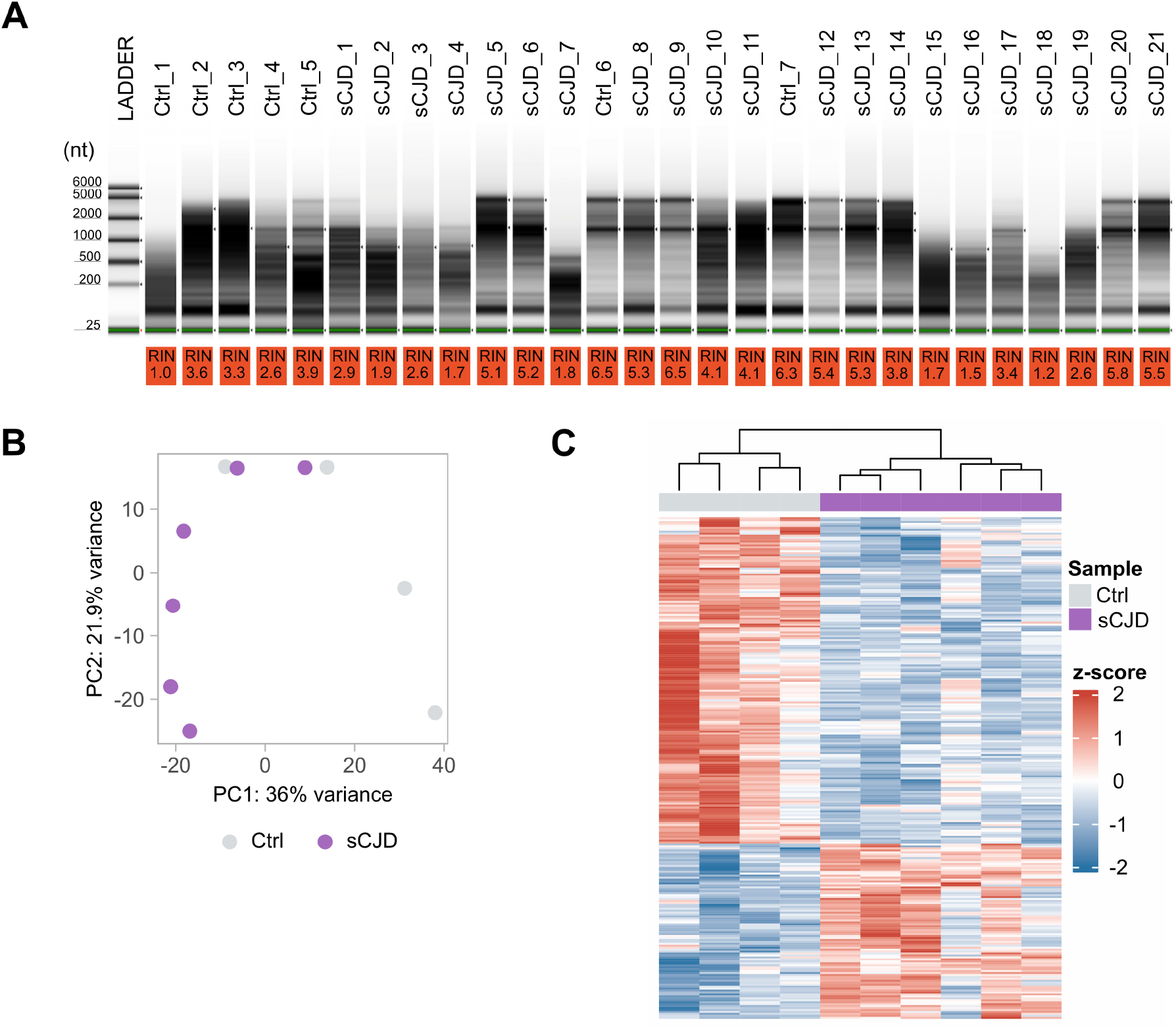
Comprehensive Assessment of Skeletal Muscle RNA Quality and Expression Patterns in sCJD Patients and Controls. **(A)** Agilent Bioanalyzer gel image depicting total RNA samples extracted from skeletal muscle tissues of both sCJD patients and control subjects. The image showcases the RNA quality assessment using the Ribosomal Integrity Number (RIN) scores, displayed alongside the respective samples. **(B)** Principal Component Analysis (PCA) plot illustrating the segregation of gene expression profiles in skeletal muscle samples between individuals with sCJD and non-sCJD controls. Each data point represents a distinct sample, with colors corresponding to two sample conditions**. (C)** Heatmap illustrating the variation in gene expression between individuals with sCJD and non-sCJD controls. Each row corresponds to a differentially expressed gene, while each column represents an individual subject from either the sCJD or control group.

**Supplementary Fig. 5.**
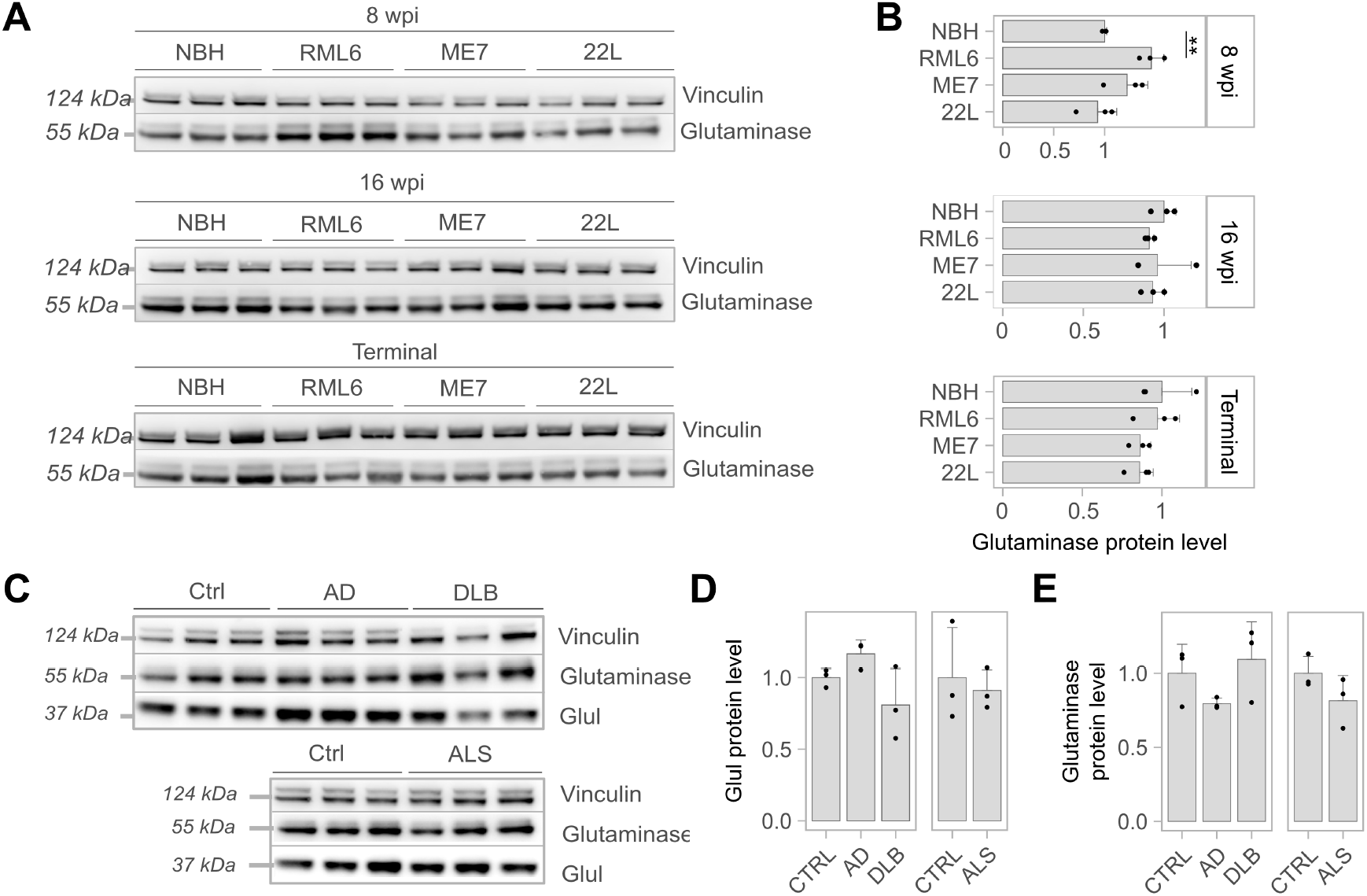
Multi-level Characterization of GLUL in Prion-Infected Mice and Neurodegenerative Disease Models, as well as Related Human Cases. (A) Western blot of Glutaminase and Vinculin protein of mice inoculated with prion strains RML6, ME7, and 22L, as well as related control (NBH). (B) Densitometry (arbitrary densitometry unit, ADU) quantification of the Western blot in Fig. 5A. In panel (C) Western blots of Glul, Glutaminase and Vinculin protein levels, of mouse models for AD, DLB and ALS, as well as related control (C57BL/6J). This Western blot presents analysis of two distinct control groups as obtained directly from different collaborators. (D) Densitometry data (ADU) quantification of Glul (normalized with related Vinculin) from the Western blot in Fig. 5C. (E) Densitometry data (ADU) quantification of Glutaminase (normalized with related Vinculin) from the Western blot in Fig. 5C. Each lane in the Western Blots represents a biological replicate. Statistical significance (*p < 0.05, **p < 0.01, ***p < 0.005, ****p < 0.001) is indicated by asterisks.

**Supplementary Fig. 6.**
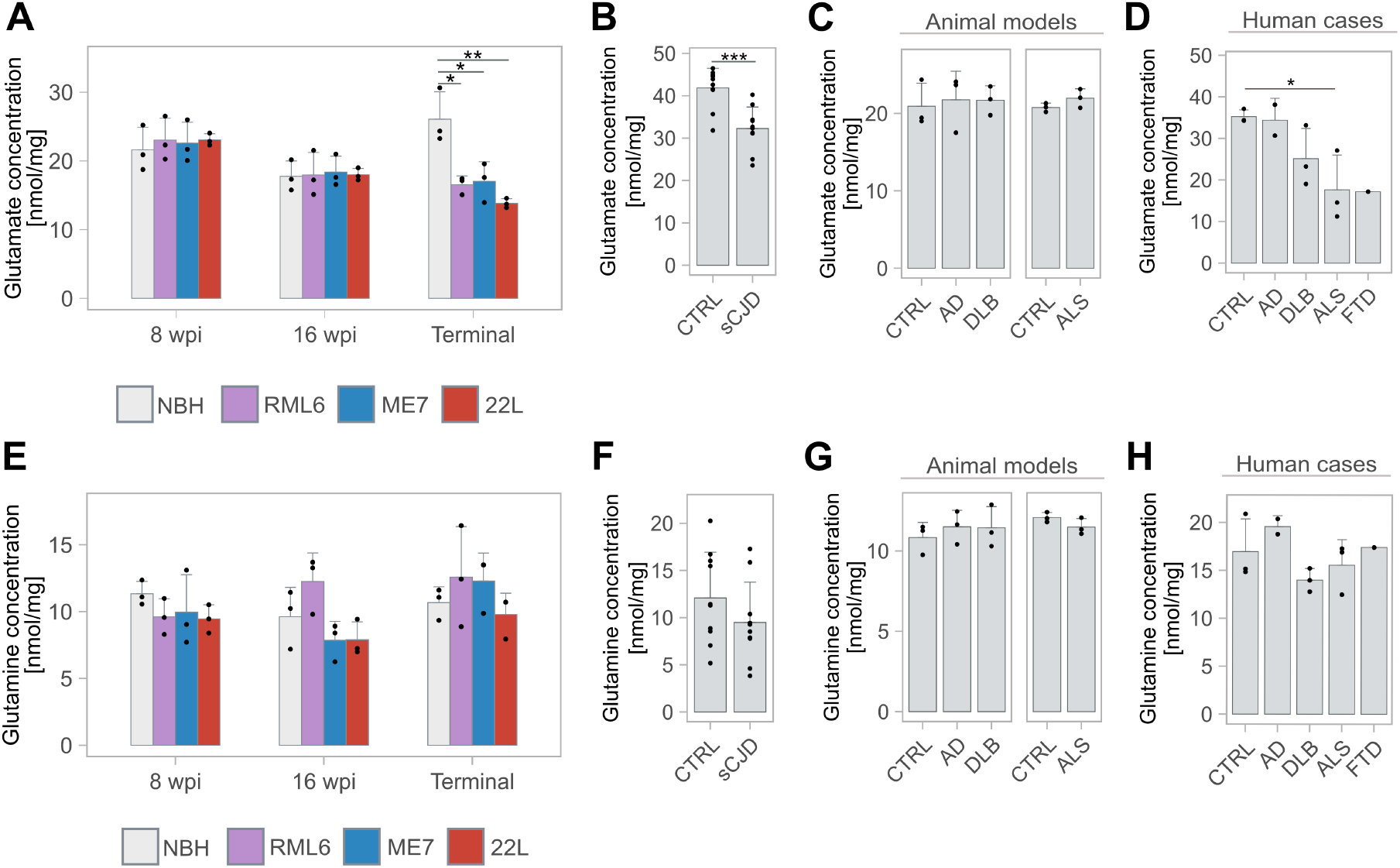
Glutamate/Glutamine Profiles in Prion-Infected Mice and Neurodegenerative Disease Models, as well as Related Human Cases. (A) Glutamate concentrations (nmol/mg) in skeletal muscle lysates of (A) mice inoculated with prion strains RML6, ME7, and 22L, as well as related control (NBH) at different timepoints, (B) of sCJD and non-sCJD control patients (C) mouse models of AD, DLB and ALS, as well as related control (C57BL/6J) and (D) human cases of AD, DLB, ALS and FTD. Glutamine concentrations (nmol/mg) in skeletal muscle lysates of (E) mice inoculated with prion strains RML6, ME7, and 22L, as well as related control (NBH) at different timepoints, (F) of sCJD and non-sCJD control patients (G) mouse models of AD, DLB and ALS, as well as related control (C57BL/6J) and (H) human cases of AD, DLB, ALS and FTD. Each dot in the graphs represents a biological replicate. Statistical significance (*p < 0.05, **p < 0.01, ***p < 0.005, ****p < 0.001) is indicated by asterisks.

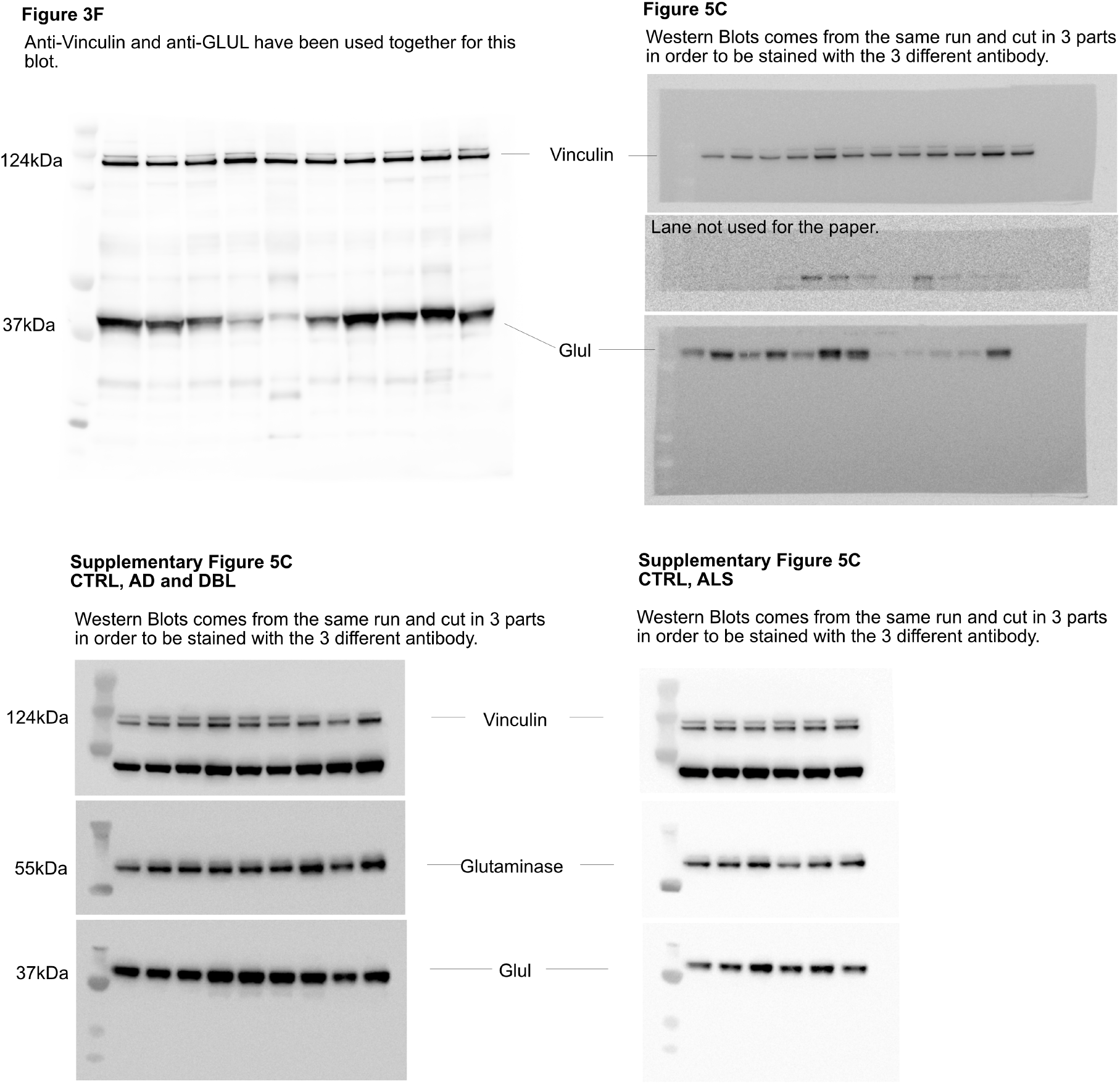

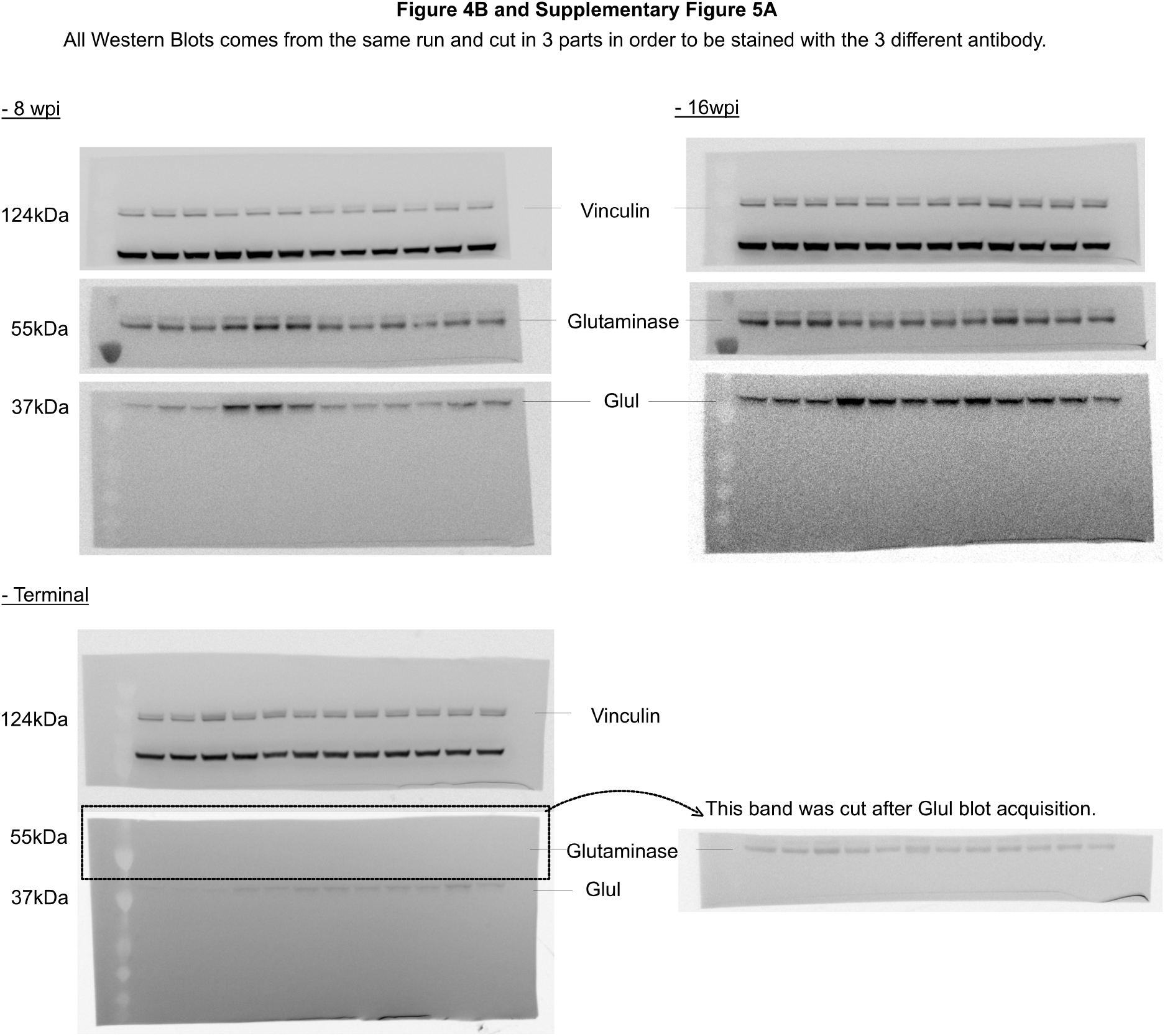

